# QT-AMP: Quanti-Tray-based amplicon sequencing for simultaneous quantification and identification of enterococci for microbial source tracking

**DOI:** 10.1101/2021.08.02.454799

**Authors:** Hidetoshi Urakawa, Taylor L. Hancock, Michael A. Kratz, Rick A. Armstrong

**Affiliations:** Department of Marine and Ecological Sciences, Florida Gulf Coast University, Fort Myers, Florida, USA; School of Geosciences, University of South Florida, Tampa, FL, 33620, USA; Lee County Environmental Laboratory, Fort Myers, FL, 33907, USA

**Keywords:** Enterococci, Enterolert, Microbial source tracking, High-throughput, amplicon sequencing

## Abstract

*Enterococcus* is ubiquitous in human feces and has been adopted as a useful indicator of human fecal pollution in water. Although regular enterococci monitoring only examines their numbers, identification of human-specific *Enterococcus* species or genotypes could help in the discrimination of human fecal contamination from other environmental sources. We documented a new approach to characterize enterococci using an amplicon sequencing platform from Quanti Trays after following the counting of most probable numbers (MPN) of enterococci. We named this method as QT-AMP (Quanti-Tray-based amplicon sequencing). We tested surface water samples collected from three rivers in southwest Florida. We detected 11 *Enterococcus* species from 45 samples in 1.1 million sequence reads. The method detected three rare species and eight cosmopolitan species (*Enterococcus faecalis, E.faecium, E. casseliflavus, E. hirae, E. mundtii, E. gallinarum, E. avium*, and *E. durans*) which have been commonly documented in various enterococci isolation studies. It is likely that the approximate detection level of QT-AMP is four orders of magnitude higher than regular 16S rRNA gene amplicon sequencing. QT-AMP revealed that a majority of Enterolert positive signals are actually the mixture of both enterococci and other facultative aerobes and anaerobes. QT-AMP may have the potential to monitor not only enterococci but also other pathogenic bacteria commonly found in natural environments. This QT-AMP could be a powerful tool to streamline the quantification and identification of enterococci and allows us to do more accurate and efficient microbial source tracking in various water management projects and human health risk assessment.

**Highlights:**

- A selected primer set (27f-519r) can differentiate over 50 *Enterococcus* species and is suitable for Illumina amplicon sequencing.
- The median of the relative contribution of enterococci reads among total sequencing reads was 82.7% and ranged between 0% and 100%.
- Enterolert signals are most likely the mixture of enterococci and other facultative aerobes and anaerobes.
- We identified eight cosmopolitan enterococci species and three rare species.
- The median of the relative contribution of non-enterococci reads among total sequencing reads was 17.3%, respectively, and ranged between 0% and 100%.

## 1. INTRODUCTION

*Enterococcus* is a gram-positive facultative anaerobic bacteria. Because of their ubiquity in human feces and persistence in the environment, *Enterococcus* has been adopted as a useful indicator of human fecal pollution in water. Traditionally, detection of enterococci has been made by membrane filtration method (e.g., U.S. EPA method 1600 membrane filtration technique) with a combination of selective medium and incubation temperature. More recently, enterococci have also been quantified using an enzymatic reaction and the most probable number (MPN) format. Now, this semi-automated MPN method for enumeration of enterococci is standardized and commercially available as Enterolert (IDEXX, Westbrook, ME, USA), a defined substrate methodology that measures a fluorescent endpoint based on enterococci metabolizing 4- methylumbelliferone-*β*-D-glucoside. The method has been widely adopted worldwide (Abbott et al., 1998; Eckner, 1998; Ramoutar et al., 2020). The method greatly improves the standardization of the counting procedure, which may cause less contamination and counting errors, and greatly reduced labor and time (Budnick et al., 1996).

Although the method is standardized and accepted broadly, the use of enterococci as indicators of human fecal pollution or contamination could be problematic (Budnick et al., 1996). Identification of origins of enterococci in water bodies is confounded by multiple sources as well as by the many factors that influence the ultimate fate of microorganisms once they are released into the environment. Enterococci are not only found in human feces but also in both wild and domestic animal feces (Harwood et al., 2000; Layton et al., 2010), birds (Devriese et al., 1992), river beds and banks, soils (Byappanahalli and Fujioka, 2004), beach sand (Ran et al., 2013; Yamahara et al., 2007), and on aquatic and terrestrial plants (Byappanahalli et al., 2003; Imamura et al., 2011).

Due to the uncertainty about how long enterococci are persistent in nature, many researchers have focused on studying their dynamics and persistence in both freshwater and saltwater environments, which can be changed by multiple environmental factors controlled by seasons, weather, hydrology, water quality, and microbial communities (Ran et al., 2013; Yamahara et al., 2007).

Identification of human-specific enterococci species or genotypes could aid in the discrimination of human fecal contamination from other environmental sources of the organisms. However, regular enterococci monitoring only addresses their numbers (Lipp et al., 2001). In the past, isolation, purification, and characterization of *Enterococcus* strains have been used to identify this bacterium at the species level (Ferguson et al., 2013; Ran et al., 2013). As an alternative approach, a cloning method of the 16S rRNA gene has been used to characterize enterococci without cultivation efforts (Sercu et al., 2011). However, these two approaches are only possible in the microbiology laboratory and not applicable in a routine manner.

To solve this issue and quantify and identify enterococcus in a streamlined fashion, we set a goal to develop a new method named QT-AMP (Quanti-Tray-based amplicon sequencing), in which first Enterolert Quanti-Tray is used for MPN counting and then pools of positive wells are used as a source for high-throughput 16S rRNA amplicon sequencing. To develop this method, we first compared three primer sets to examine which primer pair is the best to efficiently discriminate many enterococci species. Second, we tested surface water samples collected from three rivers in southwest Florida to examine how efficiently enterococci sequences were recovered and could be used for identification.

## 2. MATERIALS AND METHODS

### 2.1. Sampling sites

Water samples were collected by 12 sampling events from three rivers at routine water monitoring sampling sites of Lee County, Florida **(Fig .1).** Five sampling sites were set in the Caloosahatchee River, three in the Estero River, and four in the Imperial River. The wet and dry season samplings were conducted in August and September 2020, and January and February 2021, respectively. The Caloosahatchee River flows west from Lake Okeechobee reaching San Carlos Bay, approximately 108 km long and 62 km^2^ of surface area. The average depth of the river is 4.3 m. To provide flood control for settlement areas, intensive dredging operations were conducted in the late 1800s. After modification, now the river is a part of the Okeechobee Waterway, a manmade waterway system of southern Florida. The river is now formed by two components, compartmentalized by Franklin Lock and Dam: a narrow freshwater upstream segment and a wide tidal zone segment (Caloosahatchee River Estuary). The Estero River is a 13.7 km stream located within the Estero Bay watershed. The Imperial River is a 12.6 km stream surrounded by residential areas and has its mouth at the north end of Fish trap Bay, near the southern end of Estero Bay. A large portion of the river is covered by riverbank forests **(Fig. 1)** and intensively used for tourism (e.g., kayaking, fishing). Both rivers are impaired by fecal coliform according to the Florida Department of Environmental Protection’s (FDEP) implementation of the Impaired Waters Rule (IWR) (Gulfbase https://www.gulfbase.org/ & CHNEP Water Atlas https://chnep.wateratlas.usf.edu/). Therefore, a high abundance of fecal indicator bacteria (FIB) is problematic in these rivers.

**Fig. 1.**
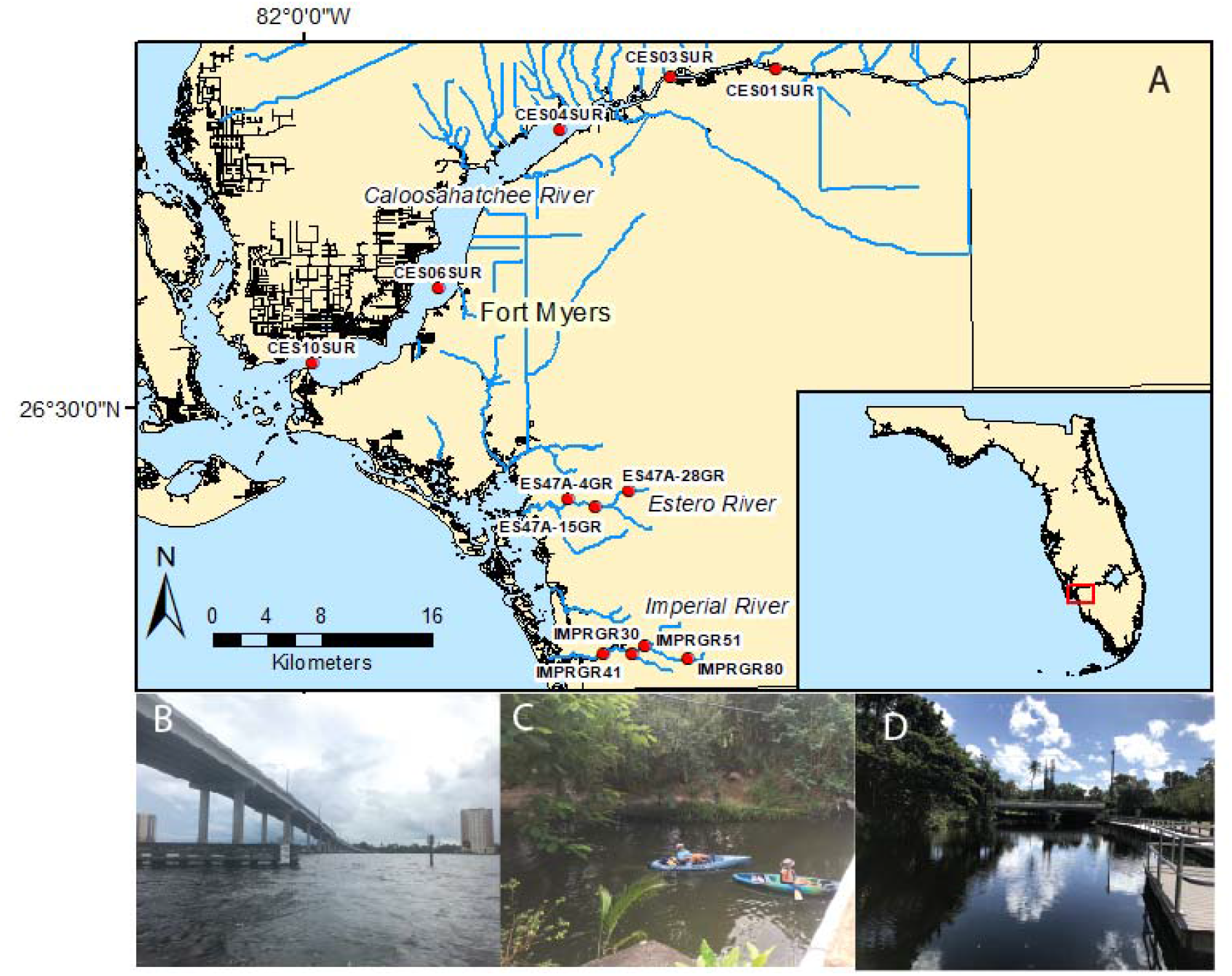
Map showing the sampling sites located in three southwest Florida rivers. Sampling map (A), the Caloosahatchee River near CES06 SUR (B), the Estero River near ES47-15GR (C), and the Imperial River near IMPRGR80 (D).

### 2.2. Water sampling

Surface water samples were collected using a clean bucket from bridges or accessible shore points. Water samples were filled in acid-washed high-density polyethylene (HDPE) bottles, kept below 8°C (iced cooler), and transported to the Lee County Environmental Lab.

### 2.3. Water quality analyses

Water quality (temperature, pH, dissolved oxygen [DO], and conductivity) analysis was conducted by a YSI multiparameter probe. Nutrients were analyzed using a SEAL AA3 autoanalyzer (Seal Analytical, Mequon, WI, USA). Total Kjeldahl nitrogen (TKN) and total phosphate (TP) were measured after heat digestion. Total nitrogen (TN) was calculated as a total of TKN and ammonia. A portion of water samples was filtered using GF/F and chlorophyll *a* was measured using a spectrophotometer after 90% acetone extraction (USEPA 445.0). Chlorophyll *a* was corrected for pheophytin. All laboratory analyses were conducted at a certificated laboratory (Lee County, Florida).

### 2.4. MPN counting using the Enterolert Quanti-Tray method

Water samples for enterococcus and *Escherichia coli* were aseptically collected and brought back to the laboratory using Thermo Scientific Security-Snap Coliform Polypropylene Water Sample Bottles with 10mg sodium thiosulfate (Thermo Fisher Scientific) and kept in ice (below 8°C). All samples were analyzed within six hours of collection following US EPA standards and the Enterolert manufacturer’s instructions. Enterolert-E ampules (IDEXX Laboratories, Westbrook, ME, USA) were aseptically added into 100 ml of water samples. The sample-reagent mixture was poured into a Quanti-Tray/2000 and sealed with the Quanti-Tray Sealer PLUS (IDEXX). The trays were incubated at 41 ± 0.5°C for 24 h. Under UV light in a dark room, yellow fluorescing positive wells were counted, and the enterococci density was calculated from the IDEXX MPN table.

### 2.5. DNA extraction from Quanti-Trays

For DNA extraction, the back of the Quanti-Tray was disinfected with 70% alcohol, and media from up to five fluorescing (positive) wells were withdrawn using sterile syringes. The withdrawn volume was recorded in each operation and approximately 10 ml of culture medium from positive wells were filtered through disposable 0.22 μm polysulfone filters (Millipore Sigma). DNA was extracted using a modified phenol-chloroform method (Urakawa et al., 2010). The purity and concentration of DNA were tested using a NanoDrop spectrophotometer (Thermo Fisher Scientific, Foster City, CA, USA) and a Qubit 2.0 Fluorometer with the Qubit dsDNA HS Assay Kit (Thermo Fisher Scientific). DNA was stored at −20°C until further analysis.

### 2.6. High-throughput DNA sequencing

A PCR primer set (27f [AGRGTTTGATCMTGGCTCAG] and 519r [GTNTTACNGCGGCKGCTG]) was used and the 27F primer contained a DNA barcode for sample identification. Samples were amplified using the HotStarTaq Plus Master Mix Kit (Qiagen, Valencia, CA, USA) with cycling conditions of 94⍰°C for 3⍰min, followed by 28 cycles of 94⍰°C for 30⍰s, 53⍰°C for 40⍰s, and 72⍰°C for 1⍰min. A final elongation step at 72⍰°C was set for 5⍰min. The success of PCR amplification was verified using agarose gel electrophoresis (2%). Samples were purified using Ampure XP beads and pooled together in equimolar concentrations. Sequencing was performed using the Illumina MiSeq platform with v3 300 base single read protocol at MR DNA (Shallowater, TX, USA). Sequence data were processed through the removal of barcodes and low-quality reads (low average quality (<Q25) or short length (<⍰200 bases), denoising, and chimera removal (Edgar et al., 2011). Forward and reverse reads were merged and operational taxonomic units (OTU) were binned by clustering sequences at 3% divergence (i.e., 97% similarity). OTUs were taxonomically classified using BLASTn against a curated database derived from GreenGenes/RDPII/NCBI. The raw sequence reads were deposited in the Short Reads Archive (SRA) and are publicly available with the accession number from SRR0000000 through SRR0000000.

### 2.7. Assessment of primer design

The best primer set was examined by in silico analysis. We collected 16S rRNA gene sequence data sets of enterococci from LPSN (List of Prokaryotic names with Standing in Nomenclature), GenBank Taxonomy, and a review manuscript (Byappanahalli et al., 2012), which were used to make the most updated species list of the genus *Enterococcus* (as of January 1, 2021). One DNA sequence was selected for one species and a total of 60 species were used for analysis **(Table S1).** The first FASTA data set was made by using 16S rRNA gene sequences provided by LPSN. Some DNA sequences originally selected were replaced by another of the same species due to relatively large numbers of undetermined nucleotide positions (*N*). Therefore, sequences containing fewer numbers of *N* and longer base pairs were selectively used for analysis. The FASTA data set were aligned by MUSCLE and manually inspected. Three sets of alignment data were created to examine the validity of the three primer sets. We assessed three primer sets: (1) 27f and 519r, (2) 515f and 806r, and (3) 515f and 926r. Sequence similarity was calculated for each species pair and a p-distance matrix was prepared for each primer set. MEGA X was used to analyze these sequence data (Kumar et al., 2018).

### 2.8. Statistical analysis

A Pearson correlation test was performed to examine the reproducibility of samples. Descriptive statistical analyses and histogram design were made using XLSTAT (Addinsoft Inc, NY, USA). A heat map was generated using Heatmapper (Babicki et al., 2016).

## 3. RESULTS

### 3.1. Which primer pair has the strongest species discrimination power?

To determine an optimum primer pair that has the strongest species discrimination power, sequence similarity was calculated among 60 *Enterococcus* species and a distance matrix was prepared for each primer set **(Table S2);** a total of 1,830 paired-distance was examined **(Table 1)**. The mean paired-distance of 27f-519r pair (8% ± 3.8% [SD]) was greater than the pairs of 515f-806r (1.3% ± 1.5%) and 515f-926r (1.3% ± 1.7%). In the case of the 27f-519r pair, a peak of relative frequency was the fourth column of the histogram (bound 0.0678-0.0904, relative frequency 0.285), while the first two columns occupied the majority of relative frequencies in the cases of 515f-806r and 515f-926r pairs **(Fig. 2).** These data suggested that the 515f-806r pair has a stronger species discrimination power than the two other primer pairs and was echoed by the DNA sequence alignment data, which contained the largest numbers of variable sites (n = 179) that aid the discrimination of *Enterococcus* species and were concentrated in the front part of the 16S rRNA gene (V1-V3 regions) **(Table 1)**.

**Fig. 2.**
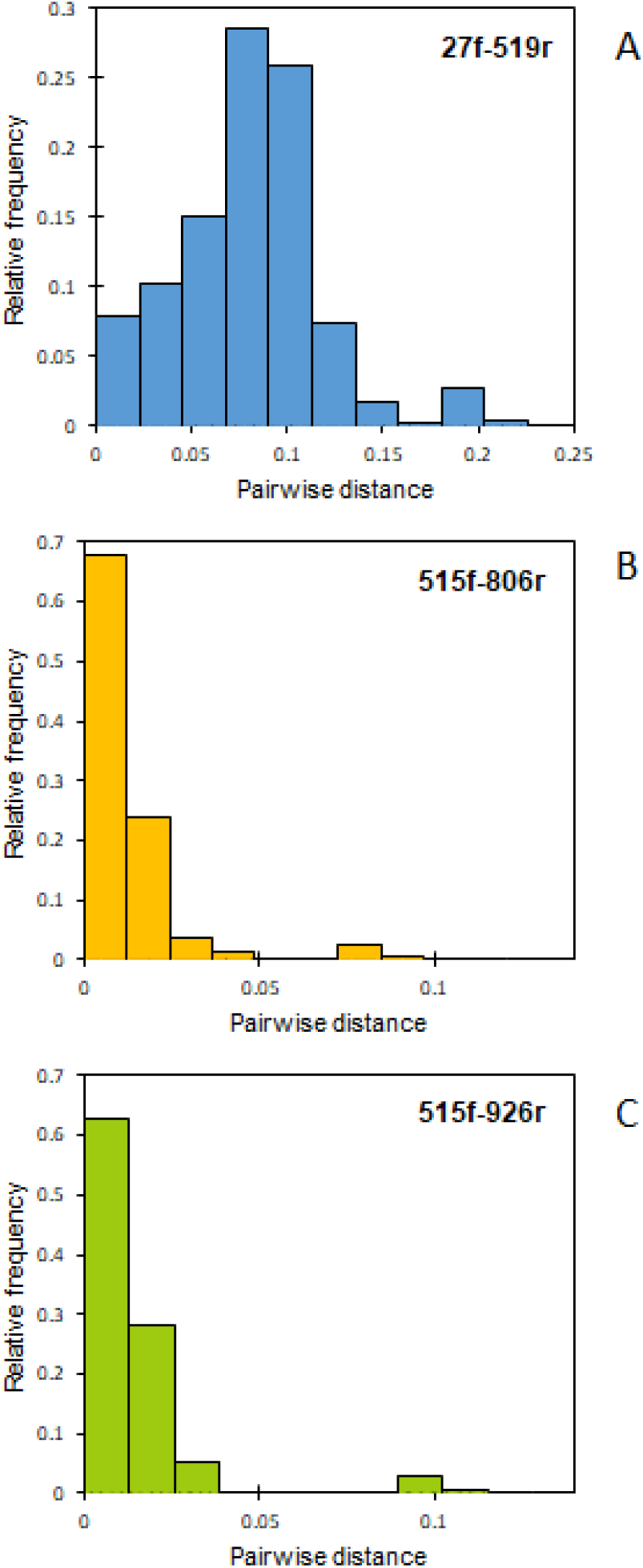
Histograms showing the relative frequency of the distribution of pairwise distance calculated from the partial 16S rRNA gene sequences of 61 *Enterococcus* species. Three primer pairs are used for each histogram. The pairwise distance of two identical sequences is zero and 5% dissimilarity is shown as 0.05.

**Table 1.** Resolution of three regions of 16S rRNA gene amplified by three primer sets

| Primer pair | Length (bp) | Variable site (n) | Covering variable region | 100% identical pairs (n) | Distinguishable species (n) | Pairwise distance |  |  |  |  |
| --- | --- | --- | --- | --- | --- | --- | --- | --- | --- | --- |
|  |  |  |  |  |  | Observation | Min | Max | Mean | SD |
| 27f & 519r | 506 | 179 | V1-V3 | 13 | 54 | 1830 | 0.000 | 0.216 | 0.080 | 0.038 |
| 515f & 806r | 253 | 37 | V4 | 225 | 25 | 1830 | 0.000 | 0.111 | 0.013 | 0.015 |
| 515f & 926r | 374 | 62 | V4-V5 | 149 | 35 | 1830 | 0.000 | 0.118 | 0.013 | 0.017 |

### 3.2. Assessment of species discrimination limitation

Our DNA database survey showed that the 16S rRNA gene sequences of 60 species of *Enterococcus* species are available (as of December 2020). All primer pairs showed that some species have 100% identities with other species, indicating that some *Enterococcus* species are not distinguishable by this high-throughput sequencing approach. We examined 100% paired match cases for each species and identified groups of *Enterococcus* clustered together. Four to five groups were found in the case of each primer pair and they consisted of 2 to 13 species **(Table S3).** Among three primer pairs, 27f-519r showed the best result and four groups found were (1) *E. caccae, E. haemoperoxidus, E. moraviensis, E. silesiacus*, and *E. ureasiticus*, (2) *E. massiliensis* and *E. songbeiensis*, (3) *E. pseudoavium* and *E. viikkiensis*, and (4) *E. porcinus* and *E. villorum*. Therefore, these listed species cannot be identified as a single species by using the 27f-519r primer pair.

### 3.3. Environmental factors and *Enterococcus* distribution

In southwest Florida, MPN counts of enterococci are routinely high and it exceeds USEPA’s enterococci criteria. The data normally show no typical point source continuously serves as a reservoir of enterococci to the river water, suggesting that the sources of enterococcus are not limited to wastewater but multiple unidentified sources **(Fig. 1).** In total, 48 water samples were collected from three rivers by 12 sampling events (six in wet and six in dry seasons) **(Table 2).** The conductivity data suggested that all three rivers were partially influenced by the mixing of seawater. The positive detection rates of enterococci in the tested water samples were 100% **(Table 3).** The geometric mean MPN counts of enterococci were highest in the Imperial River and followed by the Estero River **(Table 3).** The lowest geometric mean MPN counts were detected in the Caloosahatchee River. Over 75% of tested water samples exceeded USEPA’s statistical threshold value (STV) of 130 CFU/100 ml in the Imperial and Estero Rivers and only 5.2% in the Caloosahatchee River. Over 100% of tested water samples from each site exceeded USEPA’s geometric mean criterion of 35 CFU/100 ml in the whole sampling period in the Imperial and Estero Rivers but not from the Caloosahatchee River. Only one high number was found in the Caloosahatchee River (CES06SUR, September 2020, 782 MPN/100 ml). On that day, the water sample was characterized by low turbidity, pH, dissolved oxygen, and high nutrients, suggesting the impact of a heavy rain event **(Table 2)**. No statistically significant difference was observed for enterococci between wet and dry season samples (*p* > 0.05). The abundance of *Enterococci* and *E. coli* highly correlated in all three rivers as total (*R*^2^ = 0.83, *p* <0.001, *n* = 36) and within each river **(Table 2).**

**Table 2.**
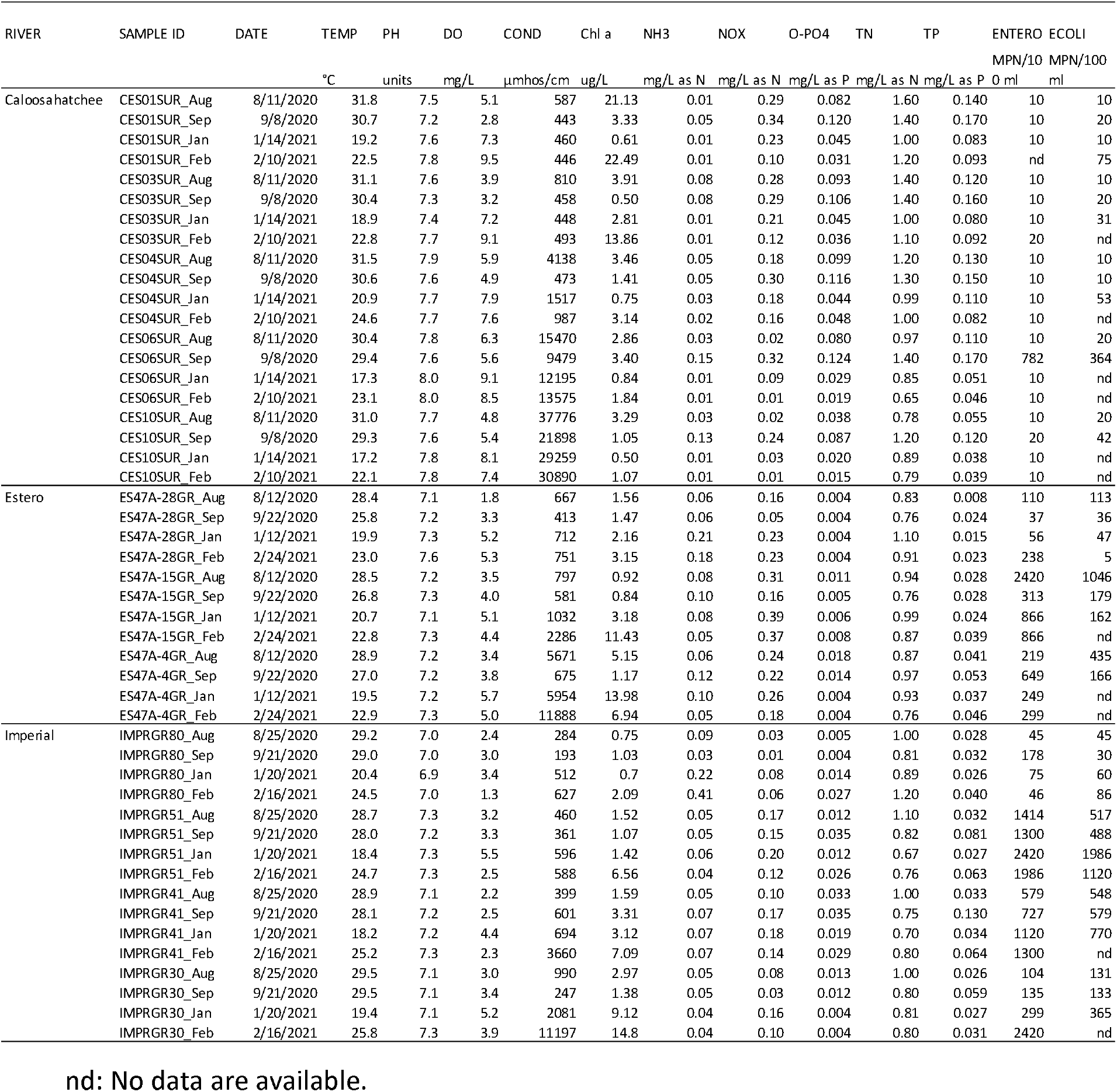
Physical and chemical parameters collected from three southwest Florida rivers

**Table 3.** Quantification of enterococci using Enterolert

| River | No of<br>sample | Sampling point | Geometric mean<br>(MPN/100 ml) | Median<br>(MPN/100 ml) | Range<br>(MPN/100 ml) | Statistical<br>threshold<br>value<br>criterion<br>(STV)<br>(frequency<br>and percent) <sup>a</sup> | Geometric mean<br>criterion<br><sup>b</sup> | Correlation with<br>*E. coli*<br>MPN<br>count<br>(R2) | Average<br>width<br>of<br>river<br>(m) <sup>c</sup> | Size of<br>the<br>watershed<br>(Km2) |
| --- | --- | --- | --- | --- | --- | --- | --- | --- | --- | --- |
| Caloosahatchee | 19 | 5 | 13.5 | 10 | 10-782 | 1/19 (5.2) | 0/5 (0) | 0.98 | 1373 | 1100 |
| Estero | 12 | 3 | 288.0 | 274 | 37-2420 | 9/12 (75) | 3/3<br>(100) | 0.79 | 17 | 931 |
| Imperial | 16 | 4 | 432.9 | 653 | 45-2420 | 12/16 (75) | 4/4<br>(100) | 0.85 | 16 | 931 |

<sup>a</sup>Tested water samples exceeded USEPA's statistical threshold value (STV) of 130 CFU/100 ml.
<sup>b</sup>Tested samples from each sampling location exceeded USEPA's geometric mean criterion of 35 CFU/100 ml in the whole sampling period.
<sup>c</sup> Average river width is determined based on the location of the sampling sites.
<sup>d</sup> The Estero River and the Imperial River share the watershed.

### 3.4. Occurrence and diversity of *Enterococcus* species

A total of 45 surface water samples containing seven duplicates were used for QT-AMP. The total and mean sequencing counts were 1,121,843 and 24,929.8 ± 1234.6 (standard error [SE], *n =* 45), respectively. The mean and median of the relative contribution of enterococci reads among total sequencing reads was 72.3 ± 4.6% (SE, *n =* 44) and 82.7%, respectively, and ranged between 0% and 100% **(Fig. 3).** A heat map was used to show the spatiotemporal distribution of enterococci species in three river water samples **(Fig. 4).** The data show a clumped distribution pattern and strong Z scores concentrated on samples from the Imperial River, which was followed by the Estero River samples. Limited enterococci species were found in the water samples from the Caloosahatchee River. This order of species distribution harmonized with the abundance of enterococci, which was measured by MPN counting **(Table 2).** In total 11 and 10 *Enterococcus* species were found from the wet and dry season samples, respectively. Unexpectedly, 10 out of 11 species overlapped each other, and *E. malodoratus* was solely found from wet season samples as a minor species **(Fig. 5).** The two highest DNA sequence reads were from *E. faecalis* (42.0% in wet season and 36.2% in dry season) and *E. casseliflavus* (34.9% in wet season and 30.1% in dry season) **(Fig. 5).** The prevalence of these two species was also shown in the heat map and these two species were major enterococci in the Caloosahatchee River **(Figs. 1 & 4).** The occurrence of each *Enterococcus* species in the water samples ranged between 100% (19/19 samples in the wet season) (*E. casseliflavus*) to 0% (0/19 samples in the dry season) (*E. malodoratus*). Many species were similarly found in both wet and dry seasons, but a few species were more frequently found in wet (*E. casseliflavus*) or dry (*E. mundtii* and *E. rivorum*) seasons **(Fig. 5).** Eleven *Enterococcus* species found in this study also well-overlapped with major *Enterococcus* species previously documented from various environments **(Table 4).**

**Fig. 3.**
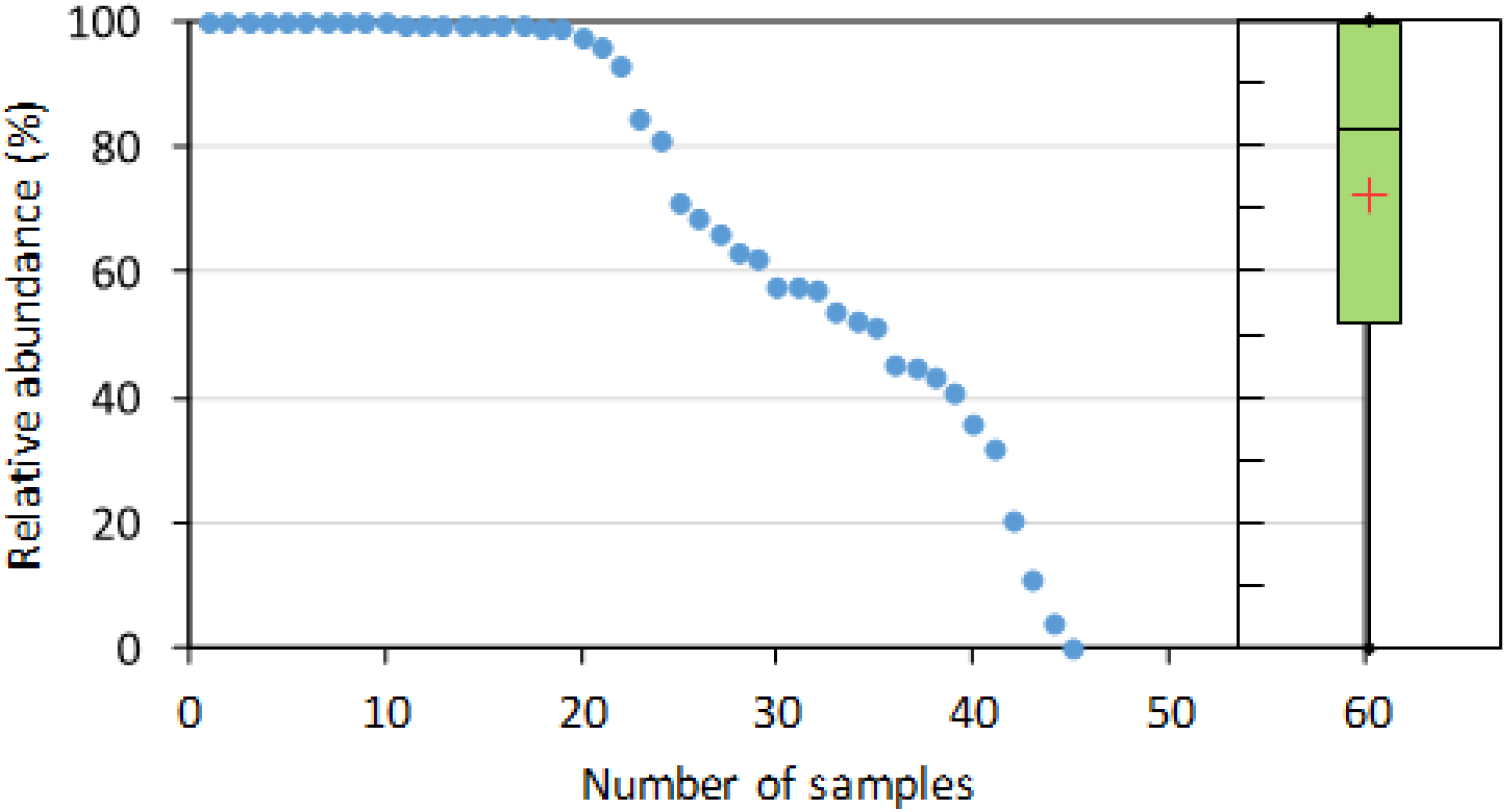
Relative abundance of enterococci in the total amplicon sequence reads. An inset box plot shows median (centerline, 82.7), first and third quartiles, mean (red cross, 72.3±4.6 [SE], *n* = 44), and range (0 to 100%).

**Fig. 4.**
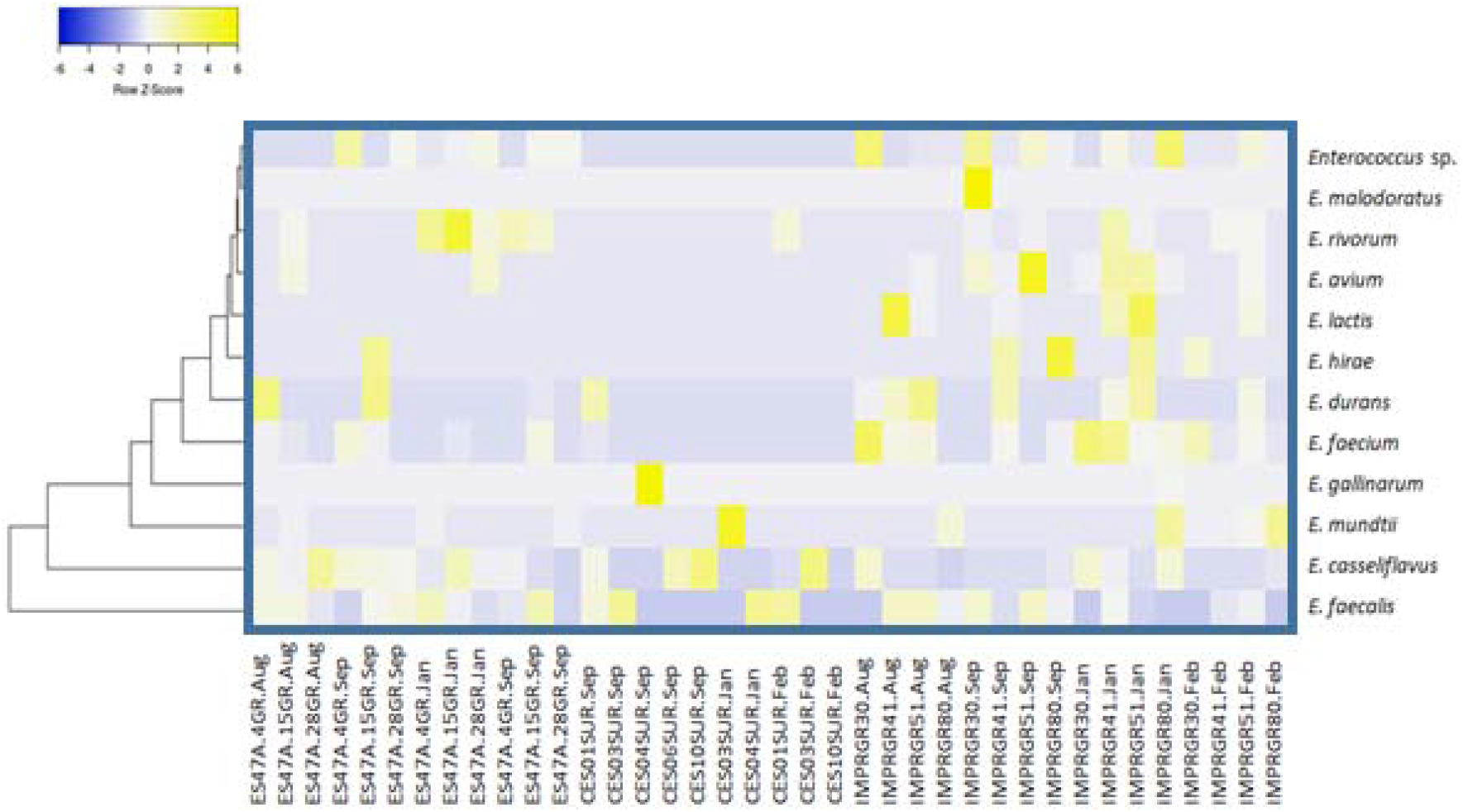
Heat map of the relative abundance of sequencing reads showing enterococci composition at the species level. Intense yellow colors indicate high standardized relative abundance values (row Z-scores), while blue colors indicate low standardized relative abundance values. Taxa are clustered using the Euclidean distance method and hierarchical clustering with the average linkage method.

**Fig. 5.**
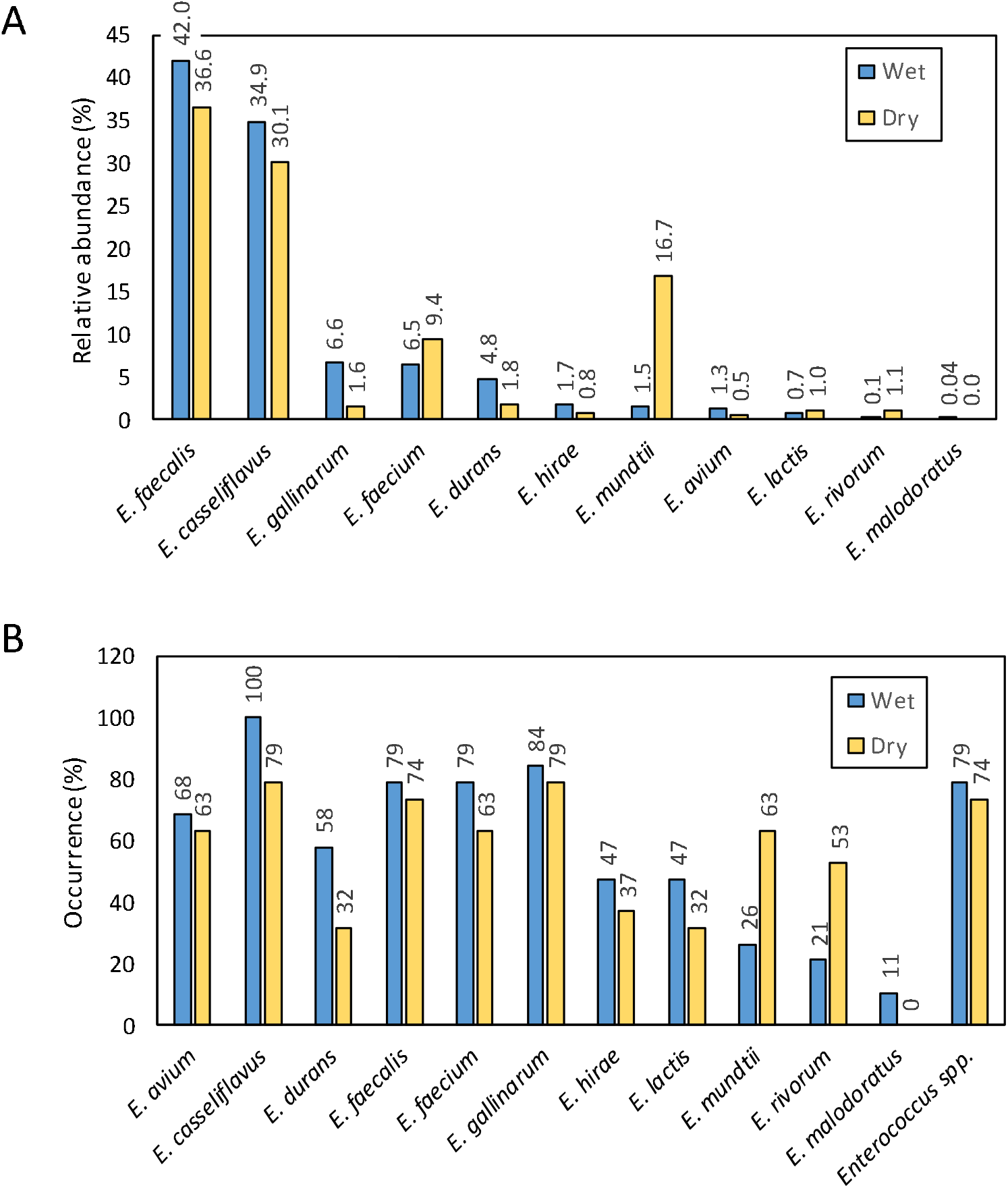
Seasonal comparison of enterococcus abundance. The read frequency of each enterococci species among the total enterococci reads (A). The occurrence of each *Enterococcus* species found among water samples (B).

**Table 4.** List of major Enterococcus species found in each study

|  | This study | Ran et al. 2013 | Ferguson et al. 2013 | Badgley et al. 2010 | Nayak et al. 2011 |
| --- | --- | --- | --- | --- | --- |
| Number of isolates |  | 2441 | 1279 | 50 | 61 |
|  | *E. casseliflavus* | *E. casseliflavus* | *E. casseliflavus* | *E. casseliflavus* | *E. casseliflavus* |
|  | *E. faecalis* | *E. faecalis* | *E. faecalis* | *E. faecalis* |  |
|  | *E. faecium* | *E. faecium* | *E. faecium* | *E. faecium* | *E. faecium* |
|  | *E. hirae* | *E. hirae* | *E. hirae* | *E. hirae* | *E. hirae* |
|  | *E. mundtii* | *E. mundtii* | *E. mundtii* | *E. mundtii* | *E. mundtii* |
|  | *E. gallinarum* | *E. gallinarum* | *E. gallinarum* |  |  |
|  | *E. avium* | *E. avium* | *E. avium* |  |  |
|  | *E. durans* | *E. durans* | *E. durans* |  |  |
|  | *E. lactis* |  |  |  |  |
|  | *E. rivorum* |  |  |  |  |
|  | *E. malodoratus* |  |  |  |  |
|  |  |  | *E. saccharolyticus* |  |  |
|  |  |  |  |  | *E. gilvus* |
|  |  |  |  |  | *E. silesiacus/moraviensis/caccae* |
| Number of species found | 11 | 8 | 9 | 6 | 6 |

### 3.5. Reproducibility of QT-AMP

Seven pairs of samples were sequenced as duplicated samples, which used two different Enterolert trays, DNA extraction, and amplicon sequencing reactions **(Fig. 6).** Four sample pairs, ES28GR-Aug, ES28GR-Sep, IMPRGR51-Jan, and IMPRGR41-Feb, showed high correlation (R^2^) among each sample and ranged between 0.893 and 0.989 (*p* <0.0001) **(Fig. 6).** One other sample (ES47A-28GR-Jan) showed a moderate correlation (0.628, *p* = 0.029). Although, two samples (CES01SUR-Sep and ES47A-28GR-Feb) showed poor reproducibility. The high reproducibility was not assured by the high number of *Enterococcus* species found in each sample and the relative abundance of *Enterococcus* sequences in total sequencing counts.

**Fig. 6.**
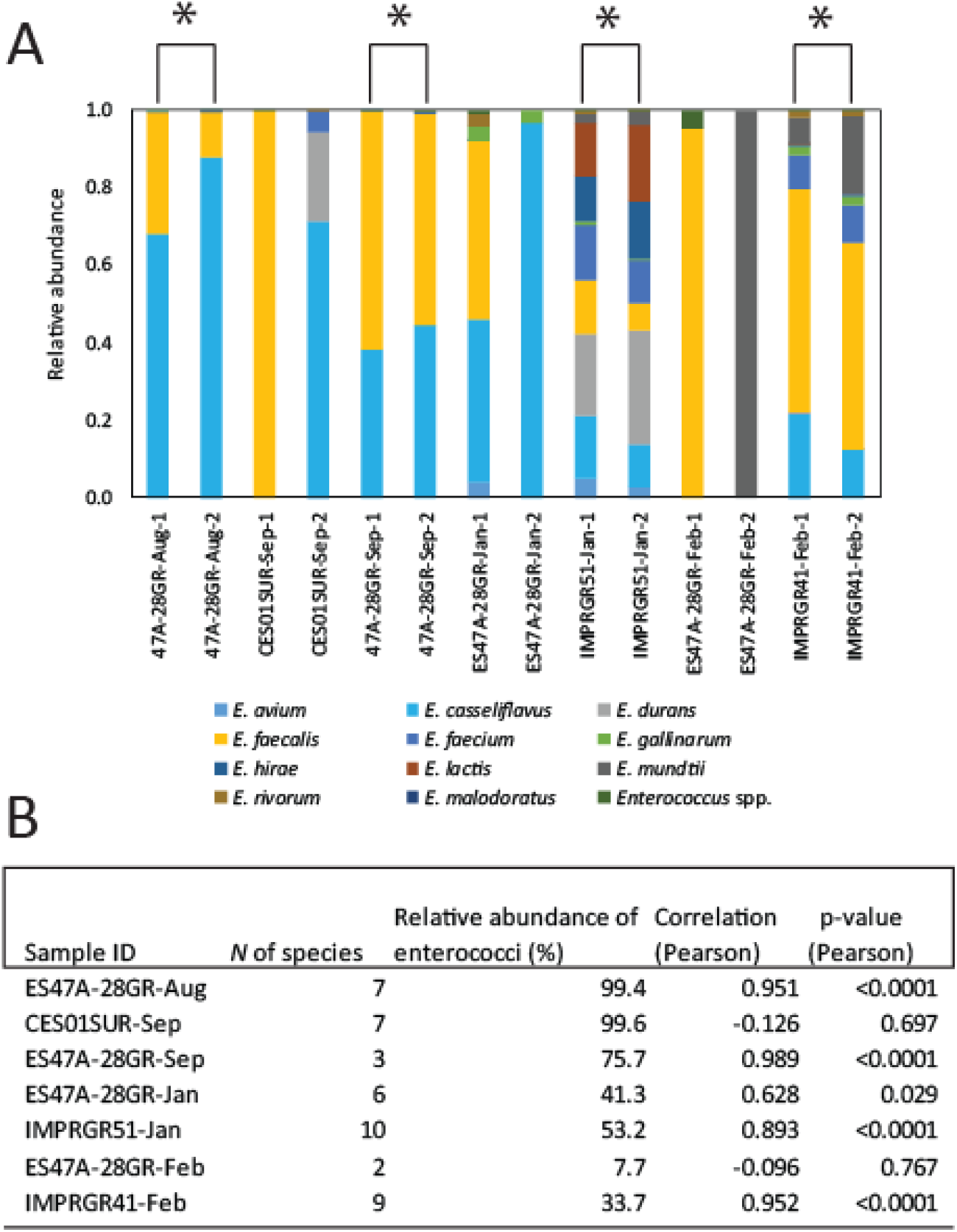
Reproducibility of seven duplicated samples tested by the relative composition of enterococci species. The relative abundance of each enterococci species (A). General statistical data and the relative abundance of enterococci among the total amplicon sequencing reads (B). (A) A star is depicted if replicates are significant (*p* < 0.01). Some species are invisible because of their minor abundance. (B) The number of enterococci species found in the averaged samples is shown.

### 3.6. Non-enterococci species found in Enterolert

Enterolert has been designed and used for quantifying enterococci in natural environments. However, it has been reported that Enterolert counting may contain non-enterococci counting (Budnick et al., 1996; Ferguson et al., 2013). In this study, we found some other non-target bacteria also grew in the wells of Enterolert. In total 16 genera in the wet season and 12 genera in the dry season were detected along with *Enterococcus* species **(Fig. 7).** As predicted, a majority of these species were facultative aerobes or anaerobes. However, a few detected bacterial taxa (*Synechococcus* and *Pelagibacter*) were not likely growing in Enterolert and at the incubation condition (no light, high nutrient, and high incubation temperature) but abundant in the original sampled water. Both *Synechococcus* and *Pelagibacter* are abundant in Caloosahatchee River water (Garcia et al. 2013).

**Fig. 7.**
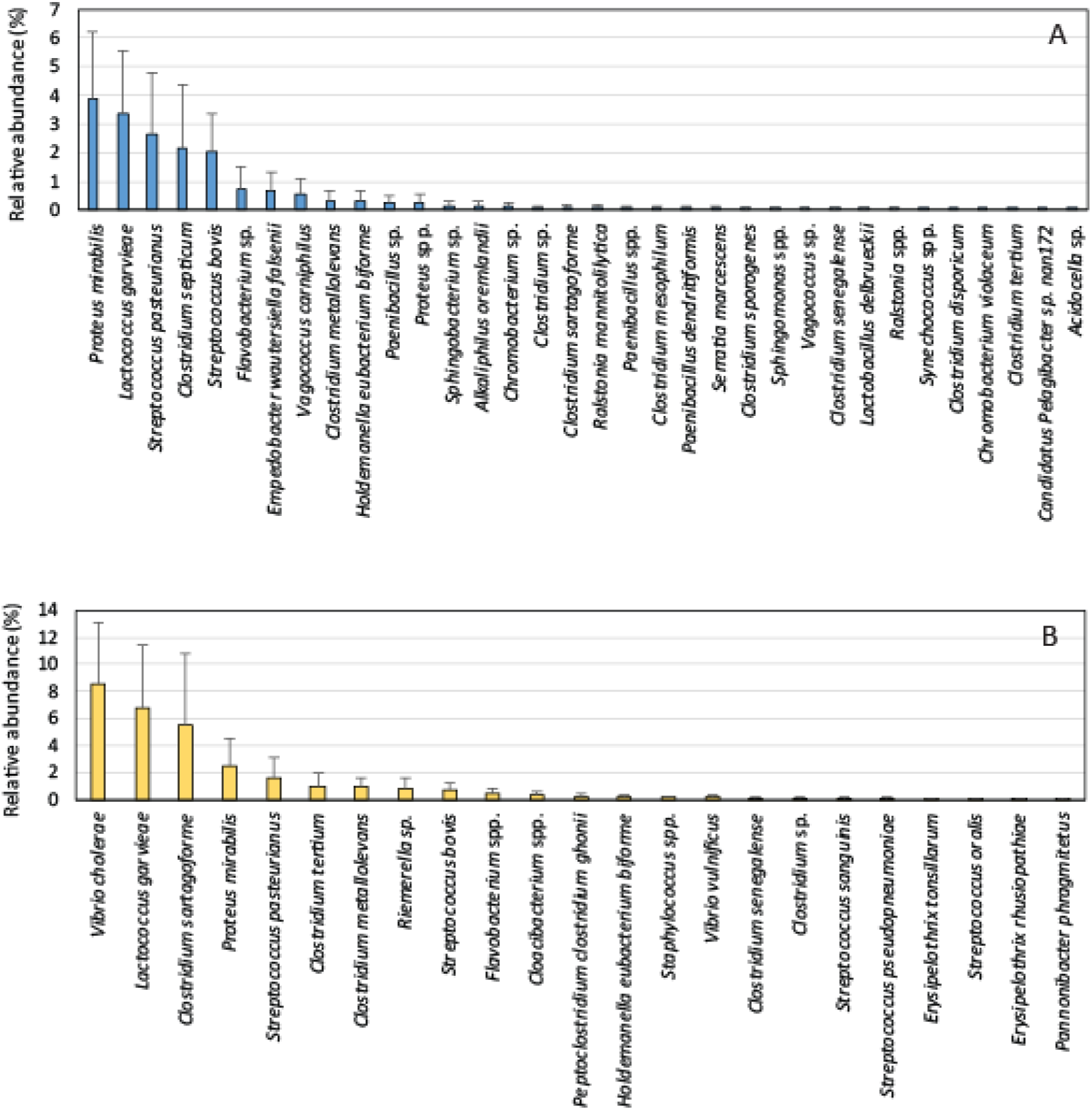
Relative abundance of detected non-enterococci bacteria. Rainy season (A) and dry season (B). Data are shown as mean and standard deviation (n = variable for each species).

## 4. DISCUSSION

Understanding the interaction between FIB density and surrounding environmental conditions has been one of the major interests of previous studies because it improves our capability to more accurately interpret the relationships between FIB densities and human health risks (Byappanahalli et al., 2012). The relationship of enterococci density and types of water bodies was previously examined in southwest Florida, and the authors concluded that the larger volumes of water allow for greater dilution and buffering against high enterococci densities; the highest densities of enterococci were found in a small stream while the lowest densities were found in Tampa Bay, which is connected to the Gulf of Mexico (Badgley et al., 2010). Our results echoed the previous study and showed that high enterococci densities were found in two small-sized rivers, the Estero River and Imperial River, while the Caloosahatchee River maintains a small number of enterococci throughout the year. Strong UV irradiation negatively influences the survival of enterococci in environments, especially in southwest Florida. Estero River and Imperial River have thick riparian vegetation that partially protects the river water from direct sunlight exposure and may prolong the survival of enterococci in river water **(Fig. 1)**.

### 4.1. *Enterococcus* as indicators of fecal contamination in recreational water

In natural water bodies, enterococci are used as a microbial indicator of environmental contamination, because they are found in high concentrations in feces, and exposure to enterococci is linked to adverse health effects in beachgoers (Lipp et al., 2001). *Enterococci* concentrations play a role in recreational water quality standards, therefore understanding of origins, survival and transport mechanisms of enterococci are necessary for effective water management. Enterolert has some advantages over the traditional counting methods and it revolutionized FIB monitoring methods (Budnick et al., 1996). In regular usage, Enterolert is designed to provide quantitative data, which is equivalent to other membrane filtration and MPN methods (Budnick et al., 1996). It does not provide qualitative data for users unless the Quanti-Tray is used to isolate and characterize enterococci after counting. Therefore, a method to allocate sources of enterococci found in surface water would be beneficial to water managers. One advantage of enterococci as FIB is that the genus contains multiple species and their habitats are diverse. A comparison of multiple studies suggests that the population structure of FIB in the environment is highly variable and the dynamics are complex (Badgley et al., 2010). QT-AMP assay is ideal for this purpose because this assay has a high sensitivity to detect various *Enterococcus* species in samples through its high-throughput format without any cultivation and identification processes. Currently, less than hundreds to up to a few thousand isolates have been phenotypically and genotypically tested (Badgley et al., 2010; Ferguson et al., 2013; Nayak et al., 2011; Ran et al., 2013), however, QT-AMP can easily provide millions of sequence reads in a short period and may be suitable to find minor species. In the present study, 11 enterococci isolates were found from water samples **(Fig. 4),** which exceeded nine species obtained from 1,279 presumptive enterococci isolates from a wide variety of environmental samples (beach water, urban runoff, and wastewater treatment plant influent & effluent) (Ferguson et al., 2013). It also exceeded eight species obtained from 2,441 *Enterococcus* strains originated from sand, sediment, water, and soil from Lake Superior. (Ran et al., 2013), suggesting the high sensitivity of QT-AMP **(Table 4).**

### 4.2. Detection of fecal indicator bacteria using high-throughput sequencing methods

One of our early challenges of the use of an amplicon sequencing technique to detect fecal indicator bacteria was the development of the method of signal amplification. For instance, in an early application of 16S rRNA amplicon sequencing to detect fecal coliform bacteria, the authors only identified a very small number of target sequences and decided to analyze the sequence data at the family level, rather than the genus or species level (Schang et al., 2016). As a part of our different projects, regular 16S rRNA gene amplicon sequence analysis was applied for water samples from the Estero River and Imperial River (unpublished data). No enterococci sequences were found in surface water samples from the Imperial River (402K in total reads from 13 samples) while 35 sequence reads from two wastewater contaminated ditch samples (all sequences were identified as *E.faecalis*) were found in the mixture of surface water, ditch water, and groundwater samples from the Estero River (623K in total reads from 31 samples). In this case, the relative abundance of enterococci found in two samples were 0.007% and 0.26% of the total reads for each sample, and 0.0056% of all sequence reads from Estero River (n = 31). In the present study, on average 72.3% of sequences (1,121K in total reads from 45 samples) were identified as enterococci. Thus, we successfully increased the detection sensitivity of enterococci targets. With the comparison of these two enterococci detection efficiency data (72.3% of QT-AMP and 0.0056% of regular amplicon sequencing), we confirmed that QT-AMP is 12,910 times more sensitive than the regular amplicon 16S rRNA gene sequencing. Therefore, we concluded that the approximate detection level of QT-AMP is four orders of magnitude higher than the regular 16S rRNA gene amplicon sequencing.

### 4.3. Non-enterococci species found in Enterolert

It has been reported that Enterolert counting likely contains some non-enterococci genera. The detection of *non-Enterococcus* species was slightly higher using Enterolert (8.4%) than for EPA Method 1600 (5.1%) when a wide variety of environmental samples (beach water, urban runoff, and wastewater treatment plant influent, and effluent) were tested (Ferguson et al., 2013). A fecal indicator bacteria study conducted in Del Rey Lagoon, California reported that 54 species were identified from 277 isolates cultured from the Quanti-Trays and many of them were not enterococci and *E. coli* (Dorsey et al., 2013). According to the U.S. EPA, the false-positive rate of enterococci isolation using mEI agar is documented as 6% (Messer and Dufour, 1998; US_Environmental_Protection_Agency, 1997). A very low level of false-positive rate (1.6%) was documented from the subtropical waters in Florida (Nayak et al., 2011). Based on QT-AMP, only one case (CES10SUR-Feb) resulted in no enterococci (1/45, 2.2%), which was a similar ratio reported from southwest Florida (Nayak et al., 2011), and only three cases (3/45, 6.7%) were conversely purely occupied by *Enterococcus* species. All other cases were a mixture of enterococci and other heterotrophic bacteria. Therefore, our data showed that a majority of Enterolert positive signals are actually the mixture of both enterococci and other heterotrophic bacteria. QT-AMP may likely have the potential to monitor not only enterococci but also other pathogenic bacteria commonly found in natural environments (Dorsey et al., 2013).

## 5. CONCLUSION

We documented a new approach to characterize enterococci using an amplicon sequencing platform from Quanti Trays after being used for counting the most probable numbers (MPN) of enterococci. We named this method as QT-AMP (Quanti-Tray-based amplicon sequencing). The importance of this work was that QT-AMP will strongly enhance semi-automated Enterolert-based enterococci monitoring methods in terms of quantity and identification. We tested surface water samples collected from three rivers in southwest Florida. We detected 11 *Enterococcus* species from 45 samples in 1.1 million sequence reads. The method detected eight cosmopolitan species (*E. faecalis, E. faecium, E. casseliflavus, E. hirae, E. mundtii, E. gallinarum, E. avium, E. durans*), which have been commonly documented from various enterococci isolation works. Thus, this identification result is compatible with previous cultivation data. The approximate detection level of QT-AMP is likely four orders of magnitude higher than the regular 16S rRNA gene amplicon sequencing. QT-AMP revealed that a majority of Enterolert positive signals are actually the mixture of both enterococci and other facultative aerobes and anaerobes. QT-AMP may likely have the potential to monitor not only enterococci but also other pathogenic bacteria commonly found in natural environments. QT-AMP has the power to streamline the quantification and identification of *Enterococcus* and can allow for more accurate and efficient microbial source tracking in various water management projects and human health risk assessment.

## Acknowledgments

We thank the Lee County Environmental laboratory for technical assistance. We also thank Drs. L. Don Duke and Serge Thomas for helpful discussion. This work was supported by the Florida Sea Grant College Program with support from the National Oceanic and Atmospheric Administration, Office of Sea Grant, U.S. Department of Commerce (NA18OAR4170085 SUB00002463).

